# Comparison of Total X-Ray Energy Absorbed by Fibroglandular Tissue and its Mass Derived from Full Field Digital Mammography and Digital Breast Tomosynthesis

**DOI:** 10.64898/2026.10.07.757391

**Authors:** John Heine, Erin Fowler, R. Jared Weinfurtner, Kathleen M. Egan

**Author notes:** Corresponding author, H. Lee Moffitt Cancer Center & Research Institute 12902 Magnolia Drive, Tampa, Florida 33612, Erin Fowler, R. Jared Weinfurtner Kathleen M. Egan.

## Abstract

This study investigated whether total x-ray energy absorbed by fibroglandular tissue and fibroglandular tissue mass derived from raw and clinical full field digital mammography (FFDM) images corresponded with the respective quantities derived from digital breast tomosynthesis (DBT). A total of 840 paired FFDM and DBT observations were analyzed separately for the craniocaudal (CC) and mediolateral oblique (MLO) orientations. Total absorbed x-ray energy and fibroglandular tissue mass were calculated using an automated breast density detection method in conjunction with the recorded average glandular dose. Relationships between FFDM and DBT quantities were evaluated in the log-log domain using linear and quadratic models, with model performance assessed by R², adjusted R², mean squared error (MSE), chi-square goodness-of-fit tests, and residual analysis. Strong associations were observed between FFDM and respective DBT quantities. For raw FFDM, linear energy models produced R² values of 0.734 and 0.901 for the CC and MLO views, respectively, while the corresponding mass models produced R² values of 0.634 and 0.870. Inclusion of a quadratic term improved model performance modestly. For clinical FFDM, linear energy models produced R² values of 0.854 and 0.919, while the corresponding mass models produced R² values of 0.721 and 0.812. The first-order coefficients for the clinical-image energy models were approximately unity (0.99–1.05), and inclusion of the quadratic term produced negligible changes in model performance and residual distributions. The goodness-of-fit tests did not reject adequate fit for any model. Clinical FFDM images can therefore provide estimates of total absorbed x-ray energy and fibroglandular tissue mass that are strongly associated with corresponding quantities derived from DBT. The approximately unit first-order coefficients and minimal benefit from quadratic modeling support a simple linear log-log calibration for clinical FFDM. This approach may facilitate combining FFDM and DBT based data in larger retrospective and longitudinal studies of cumulative mammographic x-ray exposure. These findings demonstrate proof-of-concept internal consistency but do not establish absolute dosimetric accuracy.

## Introduction

Our long-term objective is to investigate breast cancer risk associated with repeated x-ray exposures due to screening mammography. Many studies have addressed this subject and have shown the risk is small relative to the benefits provided by screening [1–3]. These studies are on firm scientific ground but relied on modeling simulations using population averages for dose, risk, and breast characteristics, rather than direct observations of radiation induced cancers. Our focus is more specific than this general case. Holding breast volume and acquisition parameters constant, a breast with a greater amount of fibroglandular tissue (quantified radiographically as breast density or dense tissue) will absorb more total x-ray energy than a more adipose breast. Breast density is an established breast cancer risk factor [4] independent of any hypothesized contribution related to an energy deposition component. Our long-term goal is to investigate whether greater cumulative x-ray energy absorbed by fibroglandular tissue during repeated screening examinations contributes to the established association between breast density and breast cancer risk. The exposure metric developed here is the energy absorbed during a given acquisition with cumulative energy due to repeated exams as the hypothesized risk-related exposure that will be investigated later. In this work, we develop suitable automated metrics for approximating the energy absorbed by the fibroglandular breast tissue volume for a mammographic acquisition. Estimates will be obtained from digital breast tomosynthesis (DBT) and full field digital mammography (FFDM) using images from women with paired acquisitions.

In the United States, the Mammography Quality Standards Act (21 CFR Part 900) establishes average glandular dose (AGD) as the standard dose metric for mammography quality control. Average glandular dose is normally expressed in millijoules per kilogram (mJ/kg), equivalent to milligray (mGy). This standardized metric is an estimate of the mean x-ray energy absorbed per unit mass of glandular tissue during a mammographic acquisition [5, 6]. In mammographic dosimetry, the glandular tissue encompasses the acinar and ductal epithelium and associated fibrous stroma, which together constitute fibroglandular tissue; stromal tissue is therefore included as a constituent of glandular tissue rather than treated as a separate component in the AGD calculation [7]. Fibroglandular tissue is radiologically dense and is distinguished from adipose tissue on mammographic images and is a significant breast cancer risk factor (i.e. breast density) [4]. Several studies have compared AGD stratified by breast density or compressed breast thickness; this work shows AGD correlated across FFDM and DBT systems, generally higher in DBT, and higher for dense breasts [8–10]. However, limited research has addressed the estimation of the total x-ray energy absorbed by the fibroglandular tissue volume or evaluated its association with breast cancer risk following many years of repeated mammographic exams. This calculation requires separating radiologically dense fibroglandular tissue from adipose tissue and estimating the corresponding fibroglandular tissue mass.

Our planned applied investigation, to study cumulative absorbed x-ray dose on breast cancer incidence, will require abstracting data from screening mammograms performed over many years. Such a study may span mammographic technologies to acquire sufficient screening histories and image data. Mammography has recently transitioned from FFDM to DBT (quasi-volumetric images reconstructed from two-dimensional projections). Using paired observations from women with both DBT and FFDM images acquired in succession, we will compare the total energy absorbed by the breast for each type of image acquisition and generate calibration curves between the two acquisition types using the paired observations. Our hypothesis is that the figures of merit (energy or mass quantities) estimated from the DBT volumes represent reference quantities for calibration, while FFDM will produce acceptable estimations permitting women with FFDM images (alone, without DBT) to be combined with those with DBT in larger studies. A previously described automated percentage of breast density (PBD) technique was used to detect mammographically dense tissue [11–14]. This primarily methodological study provides proof of concept for estimating total x-ray energy absorbed by fibroglandular tissue and establishes a practical calibration between FFDM and DBT; this may permit examinations represented by either image type to be combined in future cumulative-exposure studies.

## Methods

### Energy absorption and average glandular dose

For reference, the AGD is computed as

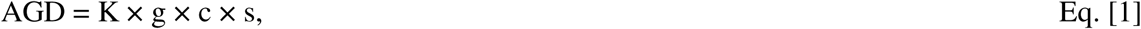

where K is the incident air kerma, g is the conversion factor from incident air kerma to AGD for a breast model with 50% glandularity, c is a correction factor for breast glandularity, s is a correction factor for the x-ray spectrum, and AGD is expressed as mJ/kg or equivalently mGy [15, 16]. At the x-ray energies used in mammography, energy absorbed by the breast, and consequently the energy represented in the AGD estimation, is primarily due to photoelectric interactions [17]. This AGD quantity is recorded in the DICOM image file (header information) from the mammography unit and is calculated using the system’s measured air kerma together with its beam calibration data and lookup tables generated with Monte Carlo simulations [15, 16]. The total x-ray energy absorbed by dense breast tissue defines our exposure metric.

### Energy absorbed by DBT acquisitions

To calculate the total energy absorbed by dense breast tissue (i.e. fibroglandular tissue) within the breast volume for a DBT acquisition, the following quantities are used: (1) n is the total number of dense voxels detected by the automated PBD method across all 2D slices, with the approximation that a pixel labeled as dense is uniform throughout the slice thickness; (2) ρ is the density of glandular tissue (1.04 g/cm³) [18]; and (3) v is the voxel volume, expressed as in-plane pixel area × 1 mm = d^2^ × 1 mm, where d is the pixel spacing in mm (recorded in the DICOM file) and the slice thickness is 1 mm with our DBT images. In DBT images, d varies across women. Making the approximation that the breast is uniformly exposed, the total energy absorbed by fibroglandular tissue is estimated as

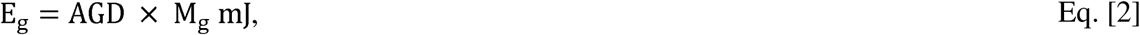

where M_g_ is the total fibroglandular mass (the subscript g is used to indicate fibroglandular throughout this work). When d is expressed in mm, the breast voxel volume is given by

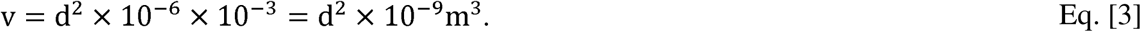

Using ρ = 1040 kg/m^3^, the total fibroglandular mass is given by

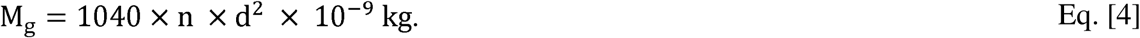

The total energy absorbed by the fibroglandular tissue volume for a single breast acquisition is then estimated as

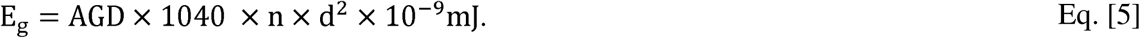

This is the total energy absorbed by fibroglandular tissue for a single breast image. For a given breast, the total energy absorbed by fibroglandular tissue is found by adding the energy from both orientations. Often images are deemed unsuitable for clinical purposes and retaken because of technical issues. For DBT and FFDM, the retake rates are about 0.38% and 0.63%, respectively [19]. Although these are relatively small proportions, an accurate accounting of the total absorbed energy over time should include these additional images. At our institution, these retakes are often available in the medical records.

### Energy Absorbed by FFDM acquisitions

We can make a similar estimation using FFDM mammograms, with an important distinction. Unlike DBT, the FFDM volume is an approximation because projected dense area is multiplied by the full compressed breast thickness at each dense pixel location. We let n equal the number of pixels in a mammogram labeled as fibroglandular tissue by the automated PBD method, t is the compressed breast thickness measured in cm, derived from the DICOM header recorded information, and d is the pixel spacing measured in mm (d is fixed for FFDM). The total dense area is then given by

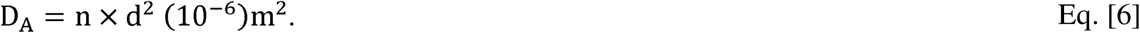

Multiplying D_A_ by the compressed breast thickness (i.e., t×10^-2^), gives an approximation of the total dense tissue volume

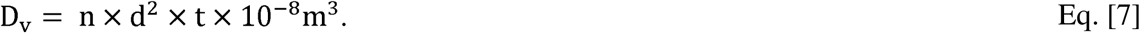

The total dense tissue mass is then given by

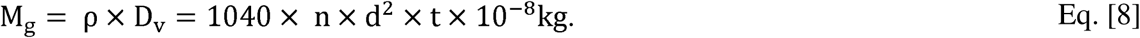

As above, the energy absorbed by fibroglandular tissue for a given image is given by

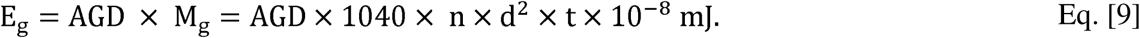

### Statistical analysis

The automated PBD method was applied to the reconstructed DBT volume slices and FFDM (raw and clinical) images to identify fibroglandular tissue. All images were acquired with Hologic Dimensions DBT units (Hologic, Inc., Marlborough, MA, USA). The 2D FFDM and DBT volume acquisitions were taken in succession (i.e., HD combination mode). Data was derived from a previous matched case-control study with 420 pairs, where the unaffected case breast and same-side control breast were investigated [11]. In total, this current study included 840 paired DBT and 2D FFDM acquisitions. In the HD-combination mode, the DBT acquisition is followed by the 2D acquisition, producing mammograms analogous to FFDM, without patient repositioning or changing the breast compression. For the 2D acquisition, both the raw and clinical FFDM images were investigated separately (i.e., modeled separately). For breast density calculations, an automated method was used to segment the breast area for both the CC and MLO orientations. For the automated PBD application, the chest wall and pectoral muscle were identified in the MLO views manually and excluded from the analysis; this area often appears over contrasted and interferes with the automated breast density detection [20]. The analysis was performed separately for the CC and MLO orientations.

To calibrate the energy between the two types of images, we assume the energy will vary over orders of magnitude when considering the variation in breast sizes and composition. Consequently, we assume a linear log-log model for paired observations

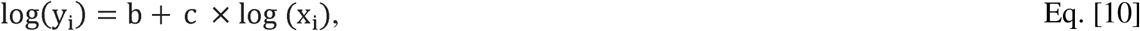

where y_i_ is the energy from DBT and x_i_ from FFDM. We refer to Eq. [10] as Model 1. Here, c will indicate if the two quantities scale linearly (c = 1), less than linearly (c < 1), or greater than linearly (c > 1), when transforming back to the measured (x_i_, y_i_) coordinates: y exp b x . Additionally, in the (x_i_, y_i_) coordinates, the transformed model approaches (0, 0) as x_i_ approaches zero. Departure from linearity in the log-log models will be evaluated by adding second and third-degree terms in log(x_i_) and jointly testing these modified models with higher-order terms against the linear log-log model.

Specifically, higher-order terms in the log-log model will be added sequentially to evaluate departures from linearity. The quadratic model is given by

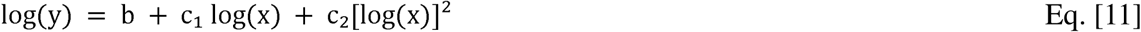

and is referenced as Model 2. If residual structure remains after including a quadratic term, a cubic term will be added accordingly. Fit will be assessed using R² and adjusted R², where adjusted R² is given by

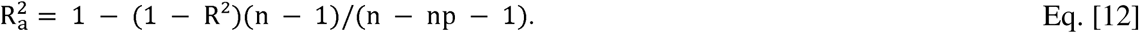

Here, R² is the coefficient of determination (the variation explained by the model), n is the number of paired observations, and np is the number of terms in the model excluding the intercept that will be used to determine whether improvements in fit obtained by adding higher-order terms justify the increased model complexity. Residual plots will also be examined for systematic departures from the fitted relationship. Models with the largest R^2^ (or R ) will be deemed appropriate in conjunction with the mean squared error (MSE) between the fitted and observed (measured) quantities. Models will also be evaluated by residual analysis, with residuals examined as a function of the fitted values to identify systematic departures from the assumed relationships. As a secondary analysis if required, the relationships in the original (x_i_, y_i_) coordinates will be fitted using third-degree polynomials, with residual analysis similarly used to assess the fidelity of these fits.

A similar analysis will be performed for dense tissue mass, with y_i_ and x_i_ representing the DBT and FFDM fibroglandular mass M_g_ estimates respectively, rather than the corresponding absorbed energy estimates. We use the same nomenclature and reference these as Model 1, Model 2, etc. This analysis will determine whether the relationship between the two image types (DBT and FFDM) is consistent for a tissue-based quantity *independent* of the absorbed-energy calculation (aside from the PBD labeling).

Because an independent physical reference standard was not available, two additional external comparisons were made with geometric estimates and values reported in previous studies. First as an order-of-magnitude energy comparison, we used a phantom-based semicircular breast model for a single CC view together with clinical mean DBT parameters, and quantities reported from other studies [7, 8]. Second, we compared our fibroglandular mass findings with other related work that used FFDM, DBT and magnetic resonance imaging (MRI) [21]. These comparisons provide the most practical external checks currently available, although they cannot replace direct validation against independently measured glandular energy deposition.

## Results

### Energy calibration

The energy calibration analysis is provided with reference to the findings presented in Table 1. The regression plots for the linear and quadratic log-log calibration models (Models 1 and 2) are shown in Figures 1 and 2, respectively, and the corresponding residual plots are shown in Figure 3. For raw FFDM CC, R^2^ increased from 0.734 to an adjusted value of 0.755, while the MSE decreased from 0.202 to 0.185, indicating a modest improvement after including the quadratic term. For clinical FFDM CC, R^2^ changed slightly (0.854 vs. 0.856), and the MSE was 0.110 vs. 0.109. For raw FFDM MLO, R^2^ increased from 0.901 to an adjusted value of 0.912, while the MSE decreased from 0.079 to 0.070, indicating a modest improvement after including the quadratic term. For clinical FFDM MLO, R^2^ remained 0.919 and the MSE 0.065 was also unchanged, indicating no improvement when adding the quadratic term. The chi-square goodness-of-fit tests did not reject the null hypothesis of adequate fit for any model (p values ranging from approximately 0.74 - 0.91). The fitted model parameters are provided in the respective captions in Figure 1 and 2, which are discussed next. For raw FFDM, the first-order coefficients were below unity in both views (0.47 for CC and 0.66 for MLO in Model 1; and 0.62 for CC and 0.72 for MLO in Model 2), while the corresponding quadratic coefficients in Model 2 were small (approximately 0.04 for both views). Importantly, for the clinical FFDM, the first order coefficient was approximately unity in both models and both views: 0.99 and 1.05 for CC and 1.0 for MLO in both models. The corresponding quadratic coefficients from Model 2 were negligible (0.03 for CC and 0.0 for MLO). The residuals from the linear and quadratic models for the clinical FFDM (or raw FFDM) were nearly indistinguishable and centered around zero, as shown in Figure 3 (Model 1 residuals are shown only). The central horizontal line that bisects the plots indicates little systematic bias. Collectively, these results demonstrate an essentially linear proportionally scaled relationship between clinical FFDM and DBT energy estimates. Because clinical images are routinely available in the archives, the findings support using the simpler linear log-log model with clinical FFDM images to estimate the corresponding DBT absorbed energy quantities. Consequently, investigating higher order log-log models with a cubic term or polynomial fitting was not required based on these findings. We note that raw FFDM images are often not archived in clinical practice [22]. These findings support using a simple linear log-log relationship to estimate the corresponding DBT energy quantities from the clinically accessible FFDM images.

**Figure 1.**
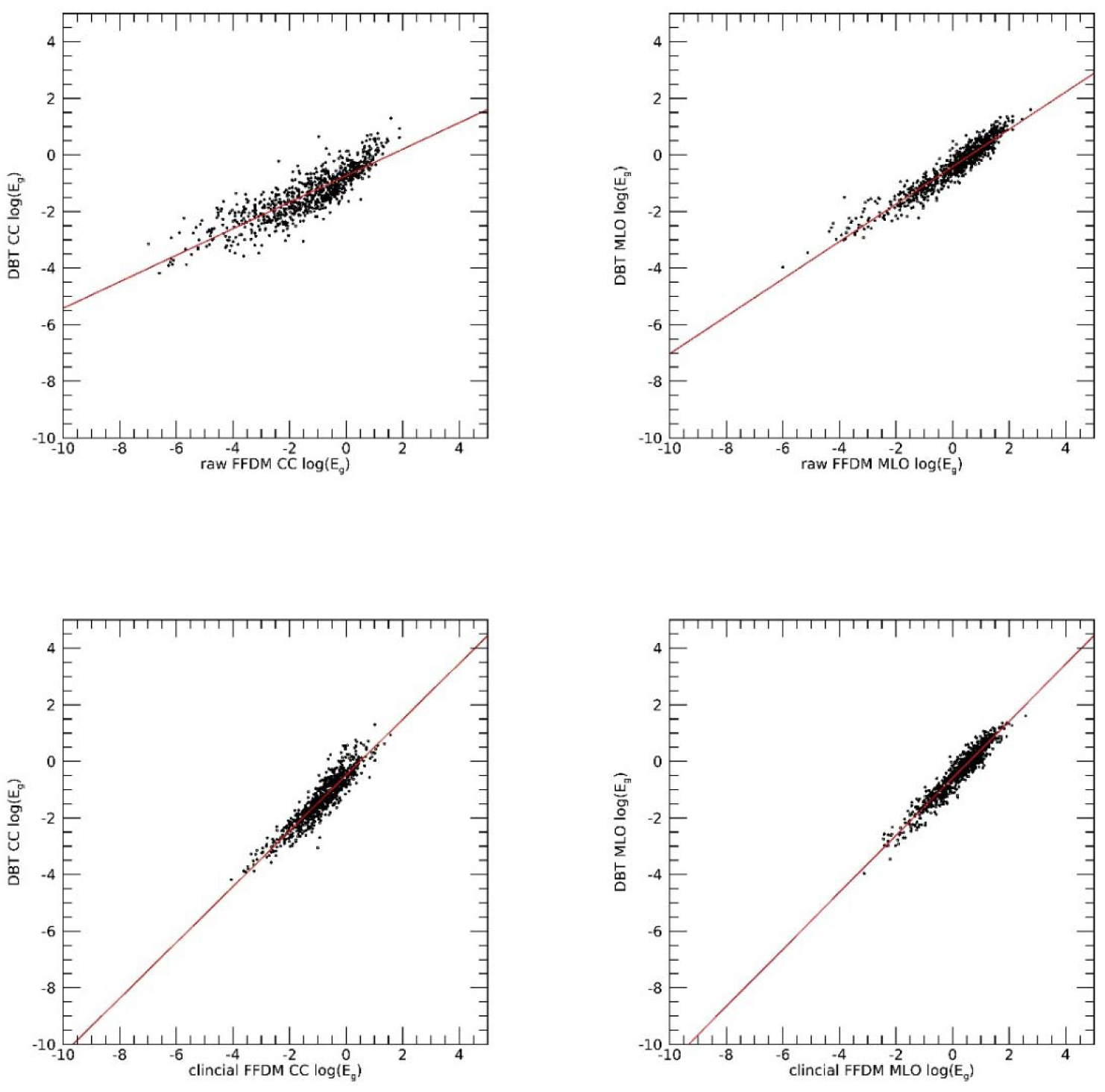
Linear Regression Analysis Model 1 Energy Relationships: The top row shows the analysis for raw FFDM images (CC on the left and MLO on the right). The bottom row shows the analysis for the clinical FFDM images. The energy (E_g_) values estimated from DBT are plotted on the vertical axes and corresponding quantities from FFDM on the horizontal axes. The regression line (solid) is compared with data (dots). The regression analysis for raw FFDM produced these parameters: (1) CC, b ≈ -0.73 and c ≈ 0.47; and (2) MLO, b ≈ -0.42 and c ≈ 0.66. Likewise, the regression analysis for the clinical FFDM produced these parameters: (1) CC, b ≈ -0.49 and c ≈ 0.99; and (2) MLO, b ≈ -0.60 and c ≈ 1.0.

**Figure 2.**
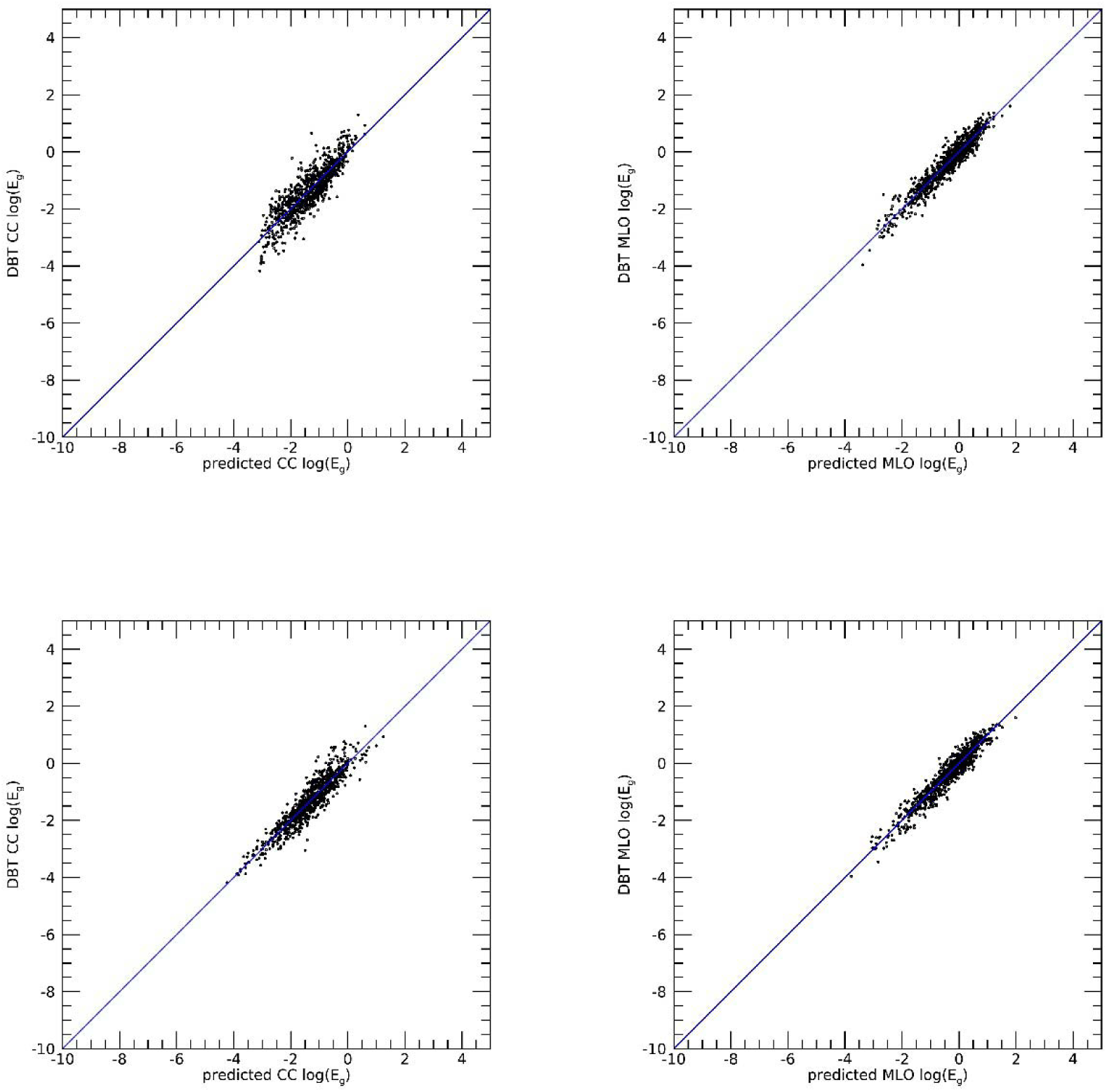
Regression Analysis for Energy Model 2: the top row shows the analysis for raw FFDM images (CC on the left and MLO on the right), and the bottom row shows the analysis for the clinical FFDM images. The energy (E_g_) values estimated from DBT are plotted on the vertical axes and fitted model values predicted from FFDM are plotted on the horizontal axes. The regression analysis for raw FFDM produced these parameters: (1) CC, b ≈ -0.70, c_1_ ≈ 0.62, and c_2_ ≈ 0.04, and (2) MLO, b ≈ -0.48, c_1_ ≈ 0.72, and c_2_ ≈ 0.04. Likewise, the regression analysis for clinical FFDM produced these parameters: (1) CC, b ≈ -0.47, c_1_ ≈ 1.05, and c_2_ ≈ 0.03; and (2) MLO, b ≈ -0.60, c_1_ = 1.0 and c_2_ ≈ 0. The diagonal line (solid) is provided in each pane for reference.

**Figure 3.**
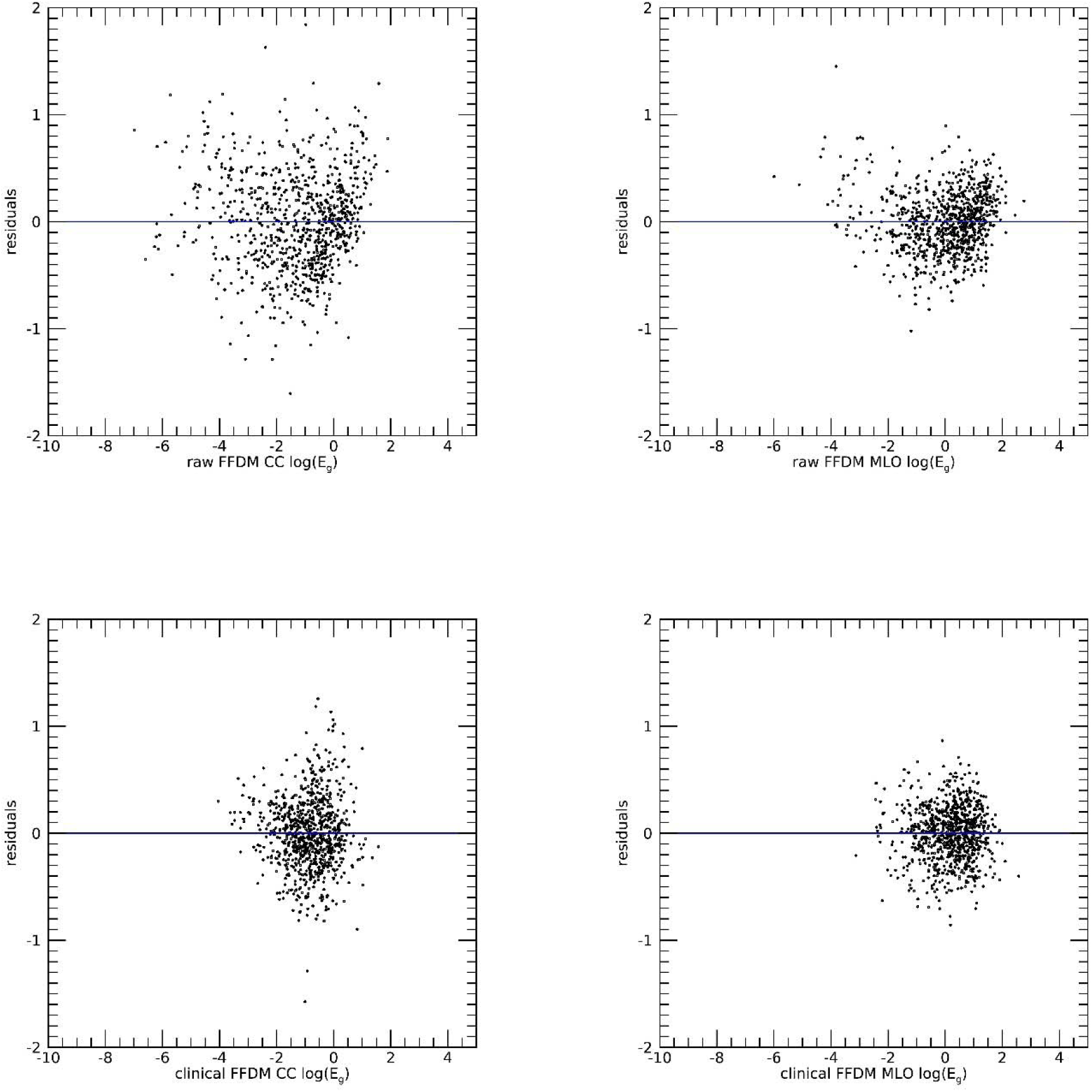
Energy Estimation Residual Analysis for Model 1: These show the residual scatter plots for energy (E_g_) modeling. The top row shows the residuals from the raw FFDM predictions and the bottom row from the clinical FFDM predictions. The solid line bisects the plane in each pane.

**Table 1.** Summary of DBT-FFDM Fibroglandular Energy and Mass Calibration Models: the linear correlation coefficient (R) and coefficient of determination (R^2^ and adjusted R ) are provided for both models. The mean squared error (MSE) is also provided for each model. Energy modeling is on the left and mass modeling on the right. Model 1 is a linear log-log model. Model 2 includes an additional quadratic log-log term.

| Model 1 (energy) | R | $R^2$ | MSE | Model 1 (mass) | R | $R^2$ | MSE |
| --- | --- | --- | --- | --- | --- | --- | --- |
| raw FFDM CC | 0.857 | 0.734 | 0.202 | raw FFDM CC | 0.796 | 0.634 | 0.144 |
| clinical FFDM CC | 0.924 | 0.854 | 0.110 | clinical FFDM CC | 0.849 | 0.721 | 0.110 |
| raw FFDM MLO | 0.949 | 0.901 | 0.079 | raw FFDM MLO | 0.932 | 0.870 | 0.043 |
| clinical FFDM MLO | 0.958 | 0.919 | 0.065 | clinical FFDM MLO | 0.901 | 0.812 | 0.062 |
| Model 2 (energy) | R | $R_a^2$ | MSE | Model 2 (mass) | R | $R_a^2$ | MSE |
| raw FFDM CC | 0.869 | 0.755 | 0.185 | raw FFDM CC | 0.815 | 0.664 | 0.132 |
| clinical FFDM CC | 0.925 | 0.856 | 0.109 | clinical FFDM CC | 0.849 | 0.721 | 0.110 |
| raw FFDM MLO | 0.955 | 0.912 | 0.070 | raw FFDM MLO | 0.940 | 0.884 | 0.039 |
| clinical FFDM MLO | 0.958 | 0.919 | 0.065 | clinical FFDM MLO | 0.904 | 0.817 | 0.061 |

### Mass calibration

The fibroglandular tissue mass calibration analysis showed a pattern, in part, similar to that of the energy comparisons. The regression plots for the linear and quadratic calibration log-log models (Models 1 and 2) are shown in Figure 4 and 5, respectively, and the corresponding residual plots are shown in Figure 6. For raw FFDM CC, R^2^ increased from 0.634 to an adjusted value of 0.664, while the MSE decreased from 0.144 to 0.132. For raw FFDM MLO, R^2^ increased from 0.870 to an adjusted value of 0.884, while the MSE decreased from 0.043 to 0.039. These findings indicate that the quadratic term improved the raw FFDM models, particularly for the MLO orientation. In contrast, only negligible changes were observed for clinical FFDM: (1) R^2^ remained unchanged 0.721 as did the MSE at 0.110 for CC; and (2) R^2^ changed from 0.812 to 0.817 and the MSE from 0.062 to 0.061 for MLO. The chi-square goodness-of-fit tests did not reject the null hypothesis of adequate fit for any mass model (p-values ranging from about 0.63 - 0.87). For the raw FFDM, the first order coefficients deviated from unity (0.40 for CC and 0.58 for MLO) in Model 1, indicating nonlinear scaling in the original coordinates, particularly for the CC orientation. For clinical FFDM, the first-order coefficients were generally close to unity, and the quadratic coefficients were relatively small, consistent with the minimal changes in R^2^, MSE, and the residual distributions (see figure captions for the regression derived parameters). Thus, because clinical images are routinely available, and the Model 1 relationship was nearly linear, the simpler linear log-log model is sufficient for estimating the corresponding DBT derived tissue mass for fibroglandular tissue.

**Figure 4.**
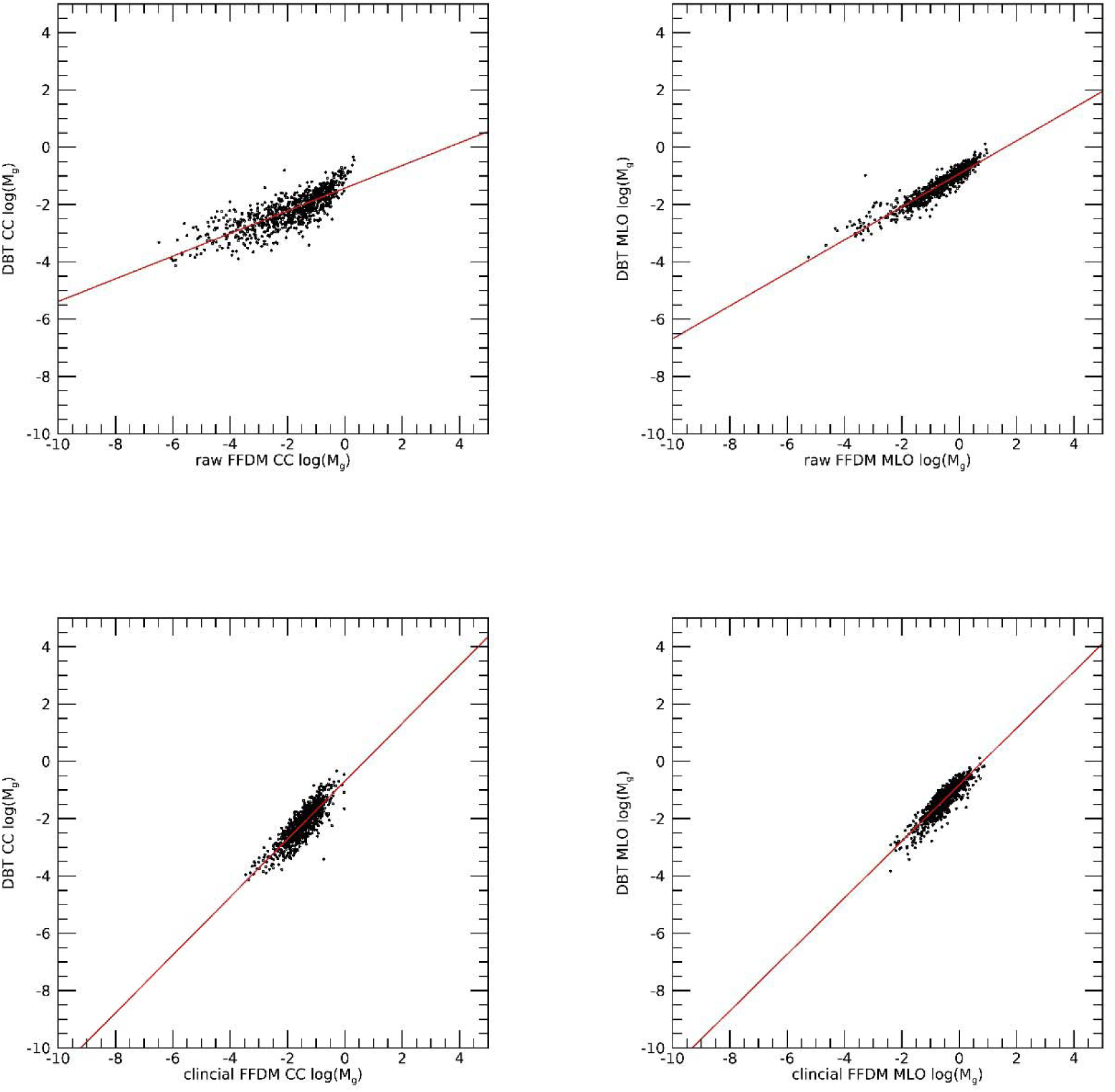
Linear Regression Analysis for Mass Model 1: the top row shows the analysis for raw FFDM images (CC on the left and MLO on the right), and the bottom row shows the analysis for the clinical FFDM images. The mass (M_g_) values estimated from DBT are plotted on the vertical axes and corresponding FFDM quantities are plotted on the horizontal axes. The regression line (solid) is compared with data (dots). The regression analysis for raw FFDM produced these parameters: (1) CC, b ≈ -1.43 and c ≈ 0.40; and (2) MLO, b ≈ -0.93 and c ≈ 0.58. Likewise, the regression analysis for clinical FFDM produced these parameters: (1) CC, b ≈ -0.70 and c ≈ 1.01; and (2) MLO, b ≈ -0.82 and c ≈ 0.98.

**Figure 5.**
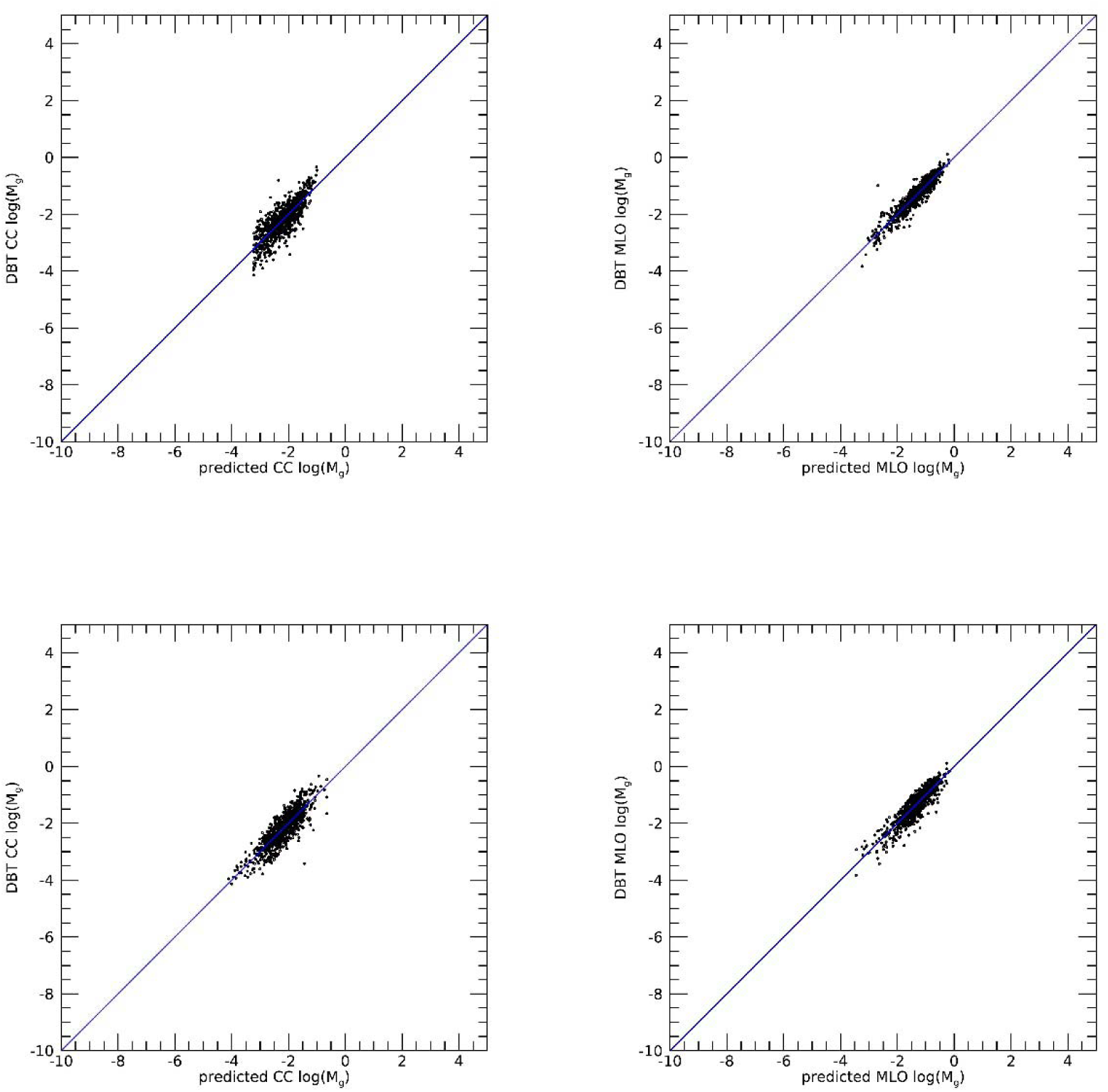
Regression Analysis for Mass Model 2: The top row shows the analysis for raw FFDM images (CC on the left and MLO on the right) and the bottom row shows the analysis for the clinical FFDM images. The mass (M_g_) values estimated from DBT are plotted on the vertical axes and fitted model values predicted from FFDM are plotted on the horizontal axes. The diagonal line (solid) is provided for reference. The regression analysis for the raw FFDM produced these parameters: (1) CC, b ≈ -1.21, c_1_ ≈ 0.66, and c_2_ ≈ 0.05 and (2) MLO, b ≈ -0.91, c_1_ ≈ 0.70, and c_2_ ≈ 0.05. Likewise, the regression analysis for clinical FFDM produced these parameters: (1) CC, b ≈ -0.63, c_1_ ≈ 1.11, and c_2_ ≈ 0.03; and (2) MLO, b ≈ -0.83, c_1_ ≈ 0.86, and c_2_ ≈ -0.10.

**Figure 6.**
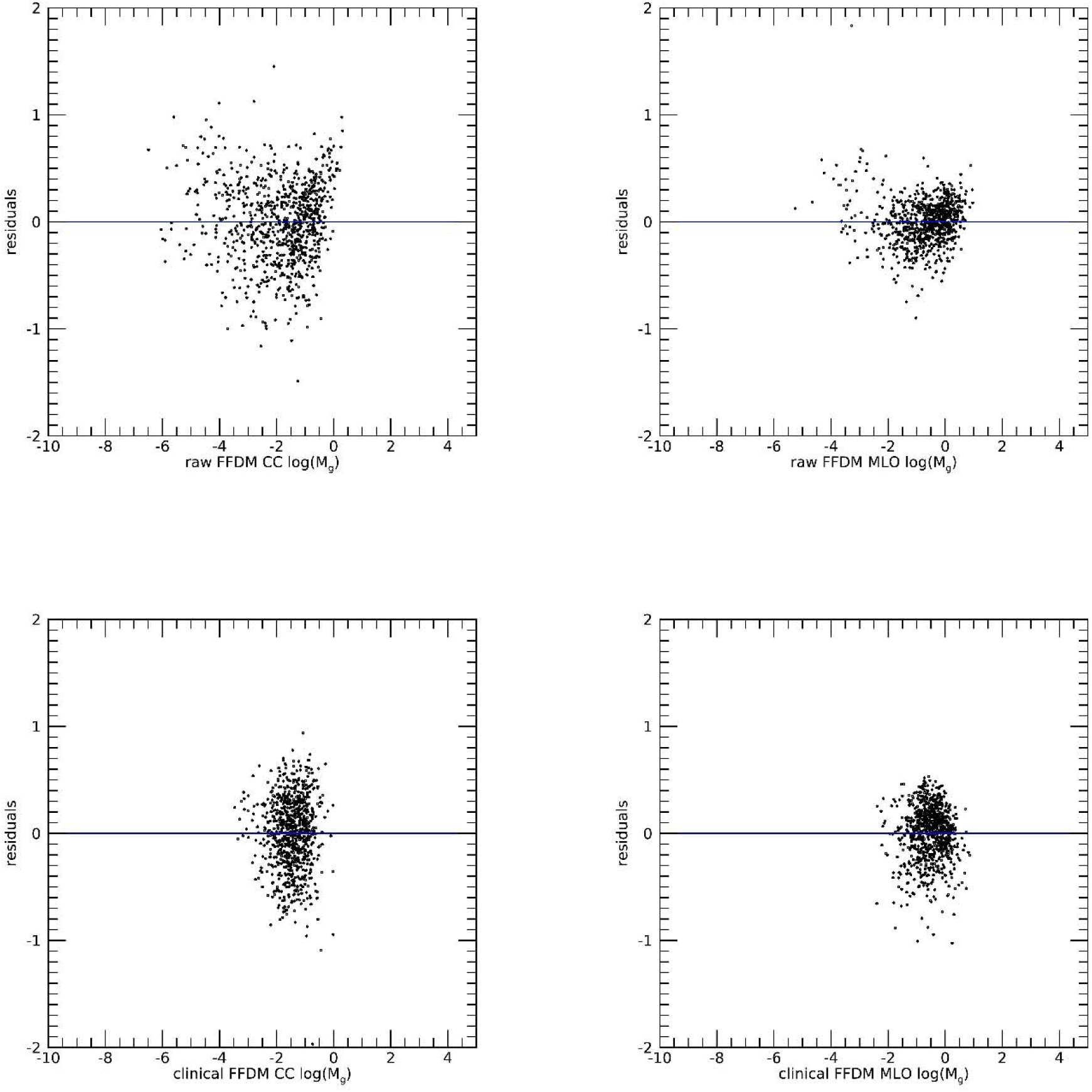
Mass Estimation Residual Analysis for Model 1: These show the residual scatter plots from the mass (M_g_) modeling. The top row shows the residuals from the raw FFDM predictions and the bottom row from the clinical FFDM predictions. The horizontal line bisects the plane in each pane.

### Energy distributions

The empirical probability distributions for E_g_ corresponding to the DBT and FFDM images are shown in Figure 7. The distribution summaries (mean, standard deviation) for the respective E_g_ quantities are provided for reference. DBT summaries (top row) were (0.37, 0.36) mJ and (0.92, 0.71) mJ for the CC and MLO mammograms, respectively. Raw FFDM summaries (middle row) were (0.65, 0.79) mJ and (1.75, 1.63) mJ for the CC and MLO mammograms, respectively. Clinical FFDM summaries were (0.57, 0.47) mJ and (1.62, 1.24) mJ for the CC and MLO mammograms, respectively. All distributions exhibited pronounced right skewness, providing justification for the logarithmic transformations used in the analysis.

**Figure 7.**
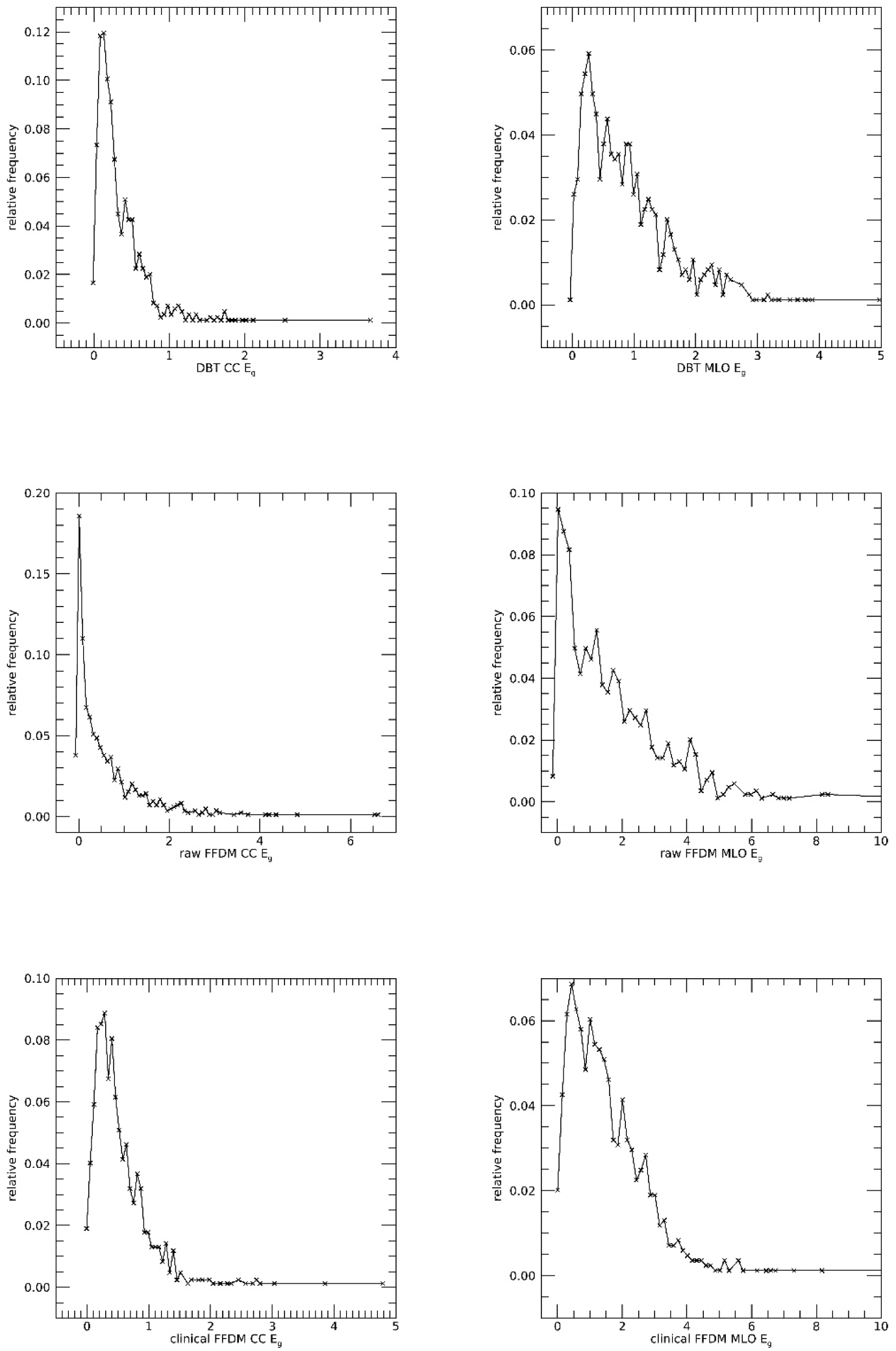
Fibroglandular Empirical Energy Probability Distribution Functions: these show the normalized histograms (empirical probability distributions) for E_g_. DBT is shown in the top row, raw FFDM in the middle row, and clinical FFDM in the bottom row. Data (x) are shown with empirical points connected with a solid line.

### External comparisons

As the first external order-of-magnitude comparison, we used a phantom-based semicircular breast model for a single CC view together with clinical mean DBT parameters. The 8.6-cm phantom radius employed by Gholamkar et al. corresponded to a projected breast area of 116.2 cm² [7]. Applying the mean compressed breast thickness of 4.8 cm reported by Gennaro et al. yielded an estimated total breast volume of 557.7 cm³ [8]. Assuming 20% - 30% glandularity and a fibroglandular tissue density of 1.04 g/cm³ (from above) resulted in an estimated fibroglandular volume of 111.5-167.3 cm³ and mass of 0.116-0.174 kg. Gennaro et al. used an automated breast-density method that operates on two-dimensional images and reported a mean CC view DBT average glandular dose of 1.858 mGy [8]. Applying this dose to the estimated fibroglandular mass yielded a total absorbed-energy range of 0.216 - 0.323 mJ for one CC acquisition of one breast, providing an approximate external assessment of the order of magnitude of the absorbed-energy estimates. These estimates are within the range of those found here. A corresponding MLO estimate was not calculated because the same geometry does not apply.

As a second external comparison, Pertuz et al. used fully automated image-analysis methods to estimate volumetric breast density and absolute fibroglandular tissue volume from FFDM, DBT, and MRI images [21]. The estimated fibroglandular tissue volumes (mean, SD) were (178, 83) cm³ for FFDM, (338, 218) cm³ for DBT, and (297, 128) cm³ for MRI. The DBT and MRI estimates were not significantly different, whereas the FFDM estimate was significantly lower; here CC and MLO views were averaged, and the DBT estimates followed from another automated breast density algorithm, whereas the FFDM estimates used a similar approach as Gennaro et al. [8]. Similar to our work, these values were algorithm-derived estimates rather than direct physical measurements of excised breast tissue. In comparison, our study produced approximately (131, 85) cm^3^ for the CC and (293,147) cm^3^ for the MLO orientations for the total fibroglandular volumes, values that are broadly similar in magnitude to those reported in this related investigation.

## Discussion and Conclusions

For the same paired observations, absorbed x-ray energy derived from DBT and FFDM showed similar correspondence with the tissue mass estimates. The consistency of these relationships across the two image acquisitions supports the internal validity of the calculations and provides complementary and sufficient proof-of-concept evidence for the present investigation, although the energy and mass estimates are not independent quantities. In future work, women with repeated and similar screening examinations acquired within a two-year window will be identified to evaluate reproducibility; reasonably close agreement is expected in the absence of substantial changes in body weight, breast composition, hormonal exposure, or acquisition conditions. The stronger associations observed for clinical FFDM may reflect the reduction of acquisition and detector-related variability through the application of vendor specific proprietary image processing, resulting in a more consistent representation of adipose and dense tissue that corresponds more closely with the reconstructed DBT measurements (also heavily processed with proprietary algorithms).

There are several qualifications of the study methods and results that warrant discussion. The study included 840 paired FFDM and DBT observations, a comparatively large sample for this type of paired-image analysis. Findings were derived from a specific population and image-acquisition setting (i.e., one vendor) and may not generalize to other populations, manufacturers, or image-processing methods. Moreover, no independent physical reference standard was available to validate the estimated fibroglandular energy and mass quantities; therefore, the agreement between FFDM and DBT derived quantities demonstrated internal consistency rather than absolute accuracy. Based on geometric modeling, our findings were consistent with external work. Model development and evaluation were also performed using the same dataset, which may produce optimistic estimates of model fit. However, the likelihood of substantial overfitting in the selected model is mitigated because it is linear, contains only one predictor, and was fitted using a relatively large number of paired observations. The nonlinear term provided negligible improvement for the routinely available clinical FFDM images and was therefore unnecessary. Nevertheless, external validation using independent populations and additional mammographic image technologies is required to establish generalizability. Although the analysis was limited to Hologic technology, Hologic systems constitute approximately 69% of the U.S. mammography installed base [23]. Thus, the findings are potentially applicable to a large proportion of mammograms acquired in current U.S. clinical practice. Consequently, more general validation is required, as even within this manufacturer the enhancement of DBT images is evolving. The MLO views were segmented using a semi-automated procedure and will require full automation in future work; notably, the resulting estimates exhibited strong monotonic relationships.

Whereas estimates of radiation-related breast cancer risk from mammographic screening have largely been based on modeling, real-world epidemiologic studies of other cancers provide evidence that radiation exposure accumulated across repeated medical imaging examinations can be associated with subsequent cancer risk. In a retrospective cohort of approximately 178,000 children and young adults who underwent CT examinations, Pearce et al. observed increasing risks of both leukemia and brain tumors with increasing cumulative radiation doses to red-bone-marrow and brain respectively; compared with patients receiving less than 5 mGy, those with a mean cumulative red-bone-marrow dose of approximately 51 mGy had a relative risk of leukemia of 3.18, while those with a mean cumulative brain dose of approximately 60 mGy had a relative risk of brain cancer of 2.82 [24]. More recently, the multinational EPI-CT cohort demonstrated an association between cumulative active-bone-marrow dose from CT examinations and hematologic malignancies, with an excess relative risk of 1.96 per 100 mGy [25]. Although these studies evaluated cumulative organ dose rather than total absorbed energy, their findings support evaluating radiation exposure across serial examinations rather than considering each examination in isolation.

Although the present analysis evaluated absorbed energy from individual observations, cumulative absorbed x-ray energy from repeated mammographic examinations may provide an additional exposure measure for investigation in relation to subsequent breast cancer incidence. The application of these calculations to serial mammograms would permit an estimation of each woman’s accumulated exposure and evaluation of its association with subsequent breast cancer risk beyond that associated with breast density, while accounting for the number and timing of screening examinations, and other established risk factors. Such an analysis would provide an important assessment of the clinical relevance of the absorbed energy measure beyond the cross-sectional measurement agreement demonstrated in the present study.

In summary, these findings demonstrate that clinical FFDM mammograms can be used to derive estimates of dense-tissue mass and total absorbed x-ray energy that are strongly associated with the corresponding quantities derived from DBT. The approximately unit first-order coefficients, negligible quadratic terms, and similar residual errors for the linear and quadratic clinical image models support use of the simpler linear log-log formulation. Because clinical FFDM images are routinely available and Hologic constitutes a large proportion of the U.S. mammography installed base, the approach may have broad practical applicability in retrospective studies encompassing past screening examinations (i.e., including mammograms that were acquired with older imaging technologies). Nevertheless, validation using independent populations and other mammographic image technologies is required. Application to serial mammograms may ultimately permit investigation of cumulative absorbed x-ray energy in relation to subsequent breast cancer risk.

## Declarations

### Author contributions

All authors contributed to the manuscript. In addition, they provided expertise in specific areas. JH, primary and corresponding author, developed the equations and code, authored the manuscript, and directed the experiments. EF is a co-author, assisted in algorithm and investigation designs, performed many of the coding tasks, applied the statistical tests, and developed the graphics in accord with the other co-authors. RW is a co-author who provided radiological and biological expertise. KE is a co-author who contributed epidemiological and biological expertise.

### Internal Review Board

All methods were carried out in accordance with relevant guidelines and regulations. All procedures were approved by the Internal Review Board (IRB) of the University of South Florida, Tampa, FL titled, Automated Quantitative Measures of Breast Density: IRB # 104715_CR000002. Mammography data was collected either (1) retrospectively on a waiver for informed consent, or (2) prospectively by gaining informed consent approved by the IRB of the University of South Florida, Tampa, FL under protocol 104715_CR000002.

### Use of Artificial Intelligence

Artificial intelligence (AI) was used to locate relevant references, which were retrieved and evaluated by the authors for relevance, and to check equations and internal reporting accuracy. The authors composed the manuscript and are responsible for its scientific content.

### Data Sharing

Image and clinical data used for this work were made public by JH. Data can be found at this reference [26]. The computer code for this work is open-source and can be found at: https://github.com/erinfowler-moffitt/Moffitt-Cancer-Center-Tampa-FL-Heine-Lab/tree/main/projects/automated-breast-density-detection-idl.

## Financial and other related disclosures

The authors have no financial or other conflicts to disclose.

